# PP2A-Cdc55 is responsible for mitotic arrest in DNA re-replicating cells in *S. cerevisiae*

**DOI:** 10.1101/2020.08.27.269696

**Authors:** Shoily Khondker, Amy E. Ikui

## Abstract

The cell cycle is an ordered process in which cells replicate their DNA in S-phase and divide them into two identical daughter cells in mitosis. DNA replication takes place only once per cell cycle to preserve genome integrity, which is tightly regulated by Cyclin Dependent Kinase (CDK). Formation of the pre-replicative complex, a platform for origin licensing, is inhibited through CDK-dependent phosphorylation. Failure of this control leads to re-licensing, re-replication and DNA damage. Eukaryotic cells have evolved surveillance mechanisms to maintain genome integrity, termed cell cycle checkpoints. It has been shown that the DNA damage checkpoint is activated upon the induction of DNA re-replication and arrests cell cycle in mitosis in *S. cerevisiae*. In this study, we show that PP2A-Cdc55 is responsible for the metaphase arrest induced by DNA re-replication, leading to dephosphorylation of APC component, Exclusion of Cdc55 from the nucleus bypassed the mitotic arrest and resulted in enhanced cell lethality in re-replicating cells. The metaphase arrest in re-replication cells was retained in the absence of Mad2, a key component of the spindle assembly checkpoint. Moreover, re-replicating cells showed the same rate of DNA damage induction in the presence or absence of Cdc55. These results indicate that PP2A-Cdc55 maintains metaphase arrest upon DNA re-replication and DNA damage through APC inhibition.

## Introduction

In order to initiate DNA replication, a pre-replicative complex (pre-RC), containing Orc1-6, Cdc6, Mcm2-7 and Cdt1, is formed on the origins of DNA replication [1]. Once the pre-RC is assembled on DNA, each component is phosphorylated by cyclin/CDK complex which causes pre-RC disassembly [2, 3]. In yeast, Orc1-6 undergoes a conformational change, Cdc6 phosphorylation is subjected to its protein degradation and Mcm2-7 is translocated to the cytoplasm after being phosphorylated [4–8]. The CDK-dependent pre-RC disassembly prohibits further steps in the initiation of DNA replication, therefore cells strictly enforce that DNA replication occurs only once in each cell cycle [9]. It has been shown that mutations in the phosphorylation sites of pre-RC components induce DNA re-replication when several mutations are combined. For example, deletion of *CDC6* N-terminus, where it is targeted by CDK (*CDC6-ΔNT*), stabilizes Cdc6. However, overexpression of this mutant alone is not sufficient to induce DNA re-replication. The combination of *GAL-CDC6ΔNT* together with an Orc6 mutant that is unable to bind to the S-phase cyclin Clb5 (*ORC6-rxl*) induces DNA damage and mitotic arrest [10, 11]. The re-replication phenotype is enhanced when additional mutations are added, such as *ORC6* phosphorylation mutant (*ORC6-ps*) or *MCM7* nuclear localization mutant (*MCM7-NLS*) [11], suggesting that these mechanisms are additive to protect cells from re-licensing and rereplication.

We have previously shown that DNA re-replication triggers DNA damage as evidenced by Ddc2 foci formation [10]. The DNA damage checkpoint is activated upon DNA re-replication in *ORC6-rxl GAL-CDC6ΔNT* cells [10, 12]. A genome wide screening identified genes involved in this process, including Mec1, Mre11-Rad50-Xrs2 (MRX complex) and Rad17-Ddc1-Mec3 (9-1-1 complex) [10]. Once DNA damage is recognized, the signaling molecules (Rad52 and Rad53) repair the damaged DNA by homologous recombination. Cell proliferation resumes once conditions are favorable again. Thus, the DNA damage signaling pathway is required for cell viability in re-replicating cells. For example, deletion of the DNA damage signaling proteins allows extensive DNA re-replication. We previously proposed a model in which forced pre-RC assembly and re-licensing will lead to DNA re-replication and DNA damage that is sensed and repaired by the DNA damage checkpoint [10].

In this study, we explore the mechanism of mitotic arrest caused by DNA re-replication in *S. cerevisiae.* Here, we show that Protein Phosphatase 2A coupled with Cdc55 regulator subunit (PP2A-Cdc55) is responsible for metaphase arrest upon induction of DNA re-replication. It is known that PP2A-Cdc55 plays a role in cell cycle progression during mitosis. Nuclear Cdc55 prevents chromosome segregation and mitotic exit through inhibition of Anaphase Promoting Complex (APC) and Cdc14 [13–15]. When Cdc55 is localized in the cytoplasm through Zds1/Zds2 interaction, PP2A-Cdc55 promotes mitotic entry by inhibiting Swe1. We used a cytoplasmic Cdc55 localization mutant, *cdc55-101,* which excludes Cdc55 from the nucleus [13, 16], to show that nuclear Cdc55 is required for cell survival of re-replicating cells. This study revealed a novel role of PP2A-Cdc55 in DNA re-replication control.

## Results

To test if PP2A^Cdc55^ is involved in the cell cycle response to DNA re-replication, we used cytoplasmic localization mutant *cdc55-101* which contains a single mutation from Gly43 to Asp [13]. *ORC6-rxl GAL-CDC6* cells with or without *cdc55-101* were serially diluted on glucose or galactose plates to test their viabilities. On galactose plates, *ORC6-rxl GAL-CDC6 cdc55-101* cells showed a more severe growth defect than the converse *CDC55* cells, indicating that there is a genetic interaction between the re-replication mutations and *CDC55* (Figure 1A, top). *ORC6-rxl,ps GAL-CDC6ΔNT* cells similarly showed a severe growth defect when combined with *cdc55-101* (Figure 1A, middle). *ORC6-rxl,ps GAL-CDC6ΔNTMCM7-NLS* cells induce extensive DNA re-replication and growth defects [10]. We observed mild synthetic lethality between *ORC6-rxl,ps GAL-CDC6ΔNTMCM7-NLS* and *cdc55-101* even on glucose plates where Cdc6 was not expressed (Figure 1A, bottom). These cells showed severe lethality on galactose plates (Figure 1A, bottom). The tetrad analysis confirmed that there is a mild synthetic lethality in *cdc55-101 ORC6-rxl,ps MCM7-NLS* cells on glucose plates (Figure 1B). These results support a role for PP2A-Cdc55 in cell survival when DNA re-replication is induced.

**Figure 1.**
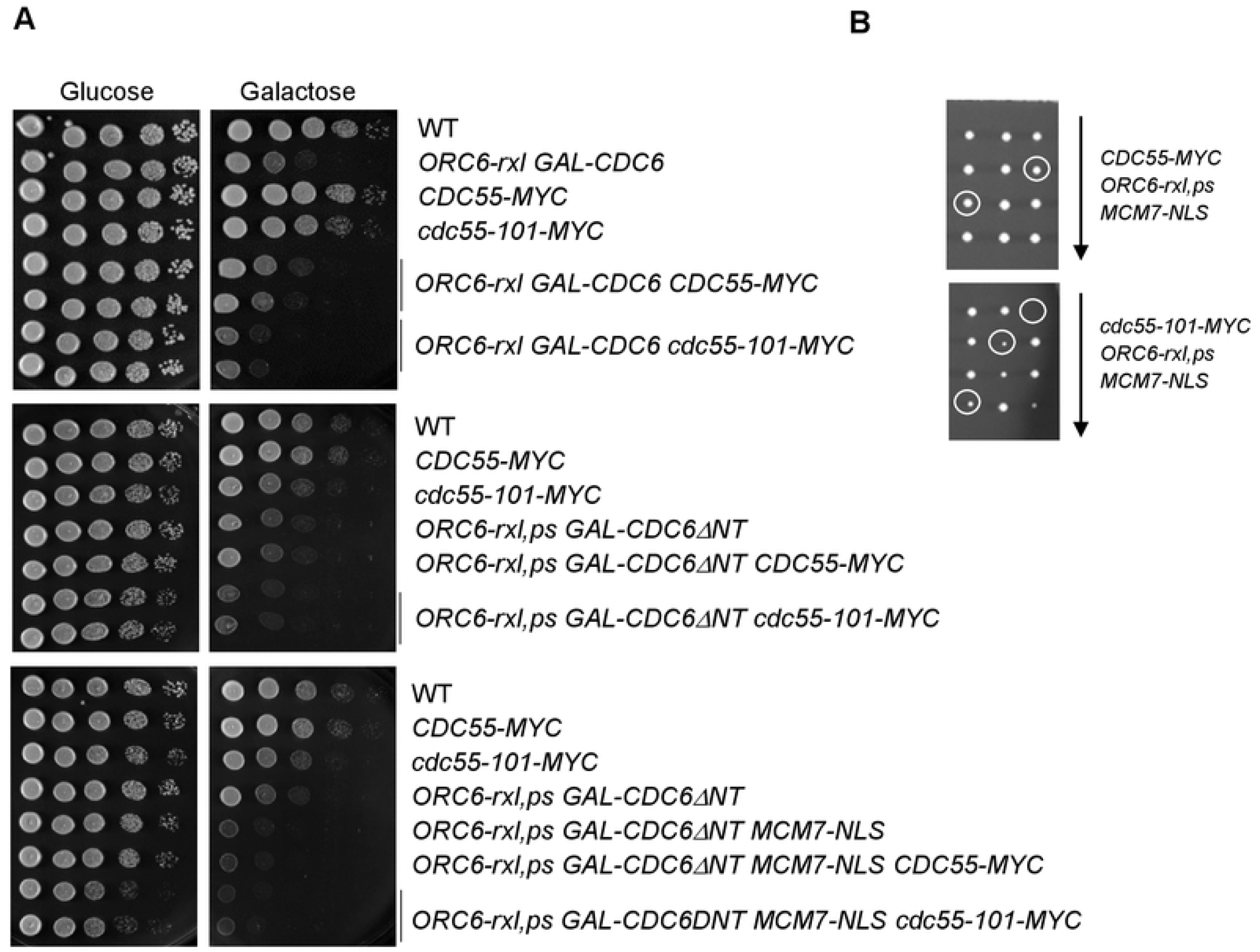
Genetic interaction between *cdc55-101* mutant and pre-RC mutants. (A) Strains with indicated genotypes were serially diluted 10-fold and plated on YEPD (glucose) or YEPG (galactose) plates. The plates were incubated at 30 degrees for 2 days. (B) Diploid cells were sporulated and the tetrads were dissected on YEPD plates. The diploid genotypes are *CDC55-MYC/CDC55 ORC6-rxl,ps/ORC6-wt MCM7-NLS/MCM7-wt* (top) or *cdc55-101-MYC/CDC55 ORC6-rxl,ps/ORC6-wt MCM7-NLS/MCM7-wt* (bottom). The circled colonies contain all three mutations.

Cells with elongated buds are reminiscent of mitotic arrest after DNA re-replication [10]. To test if PP2A-Cdc55 is involved in the mitotic arrest, the cell cycle profile was examined in DNA re-replicating cells in the presence or absence of *cdc55-101.* The majority of *ORC6-rxl,ps GAL-CDC6* cells contained 2C DNA content, indeed confirming mitotic arrest. When combined with *cdc55-101,* cell cycle profile was changed; 1C DNA with reduced 2C DNA content. It suggests that the mitotic arrest in the re-replicating cells was reversed (Figure 2A, left). Consistent with prior reports, *ORC6-rxl,ps GAL-CDC6 MCM7-NLS* cells showed tight 2C arrest (Figure 2A, right) [10]. Addition of the *cdc55-101* mutation in the *ORC6-rxl,ps GAL-CDC6 MCM7-NLS* cells showed a reduced mitotic arrest, indicating that the cells escaped from the mitotic arrest. We conclude that nuclear Cdc55 is responsible for the mitotic arrest in overreplicating cells.

**Figure 2.**
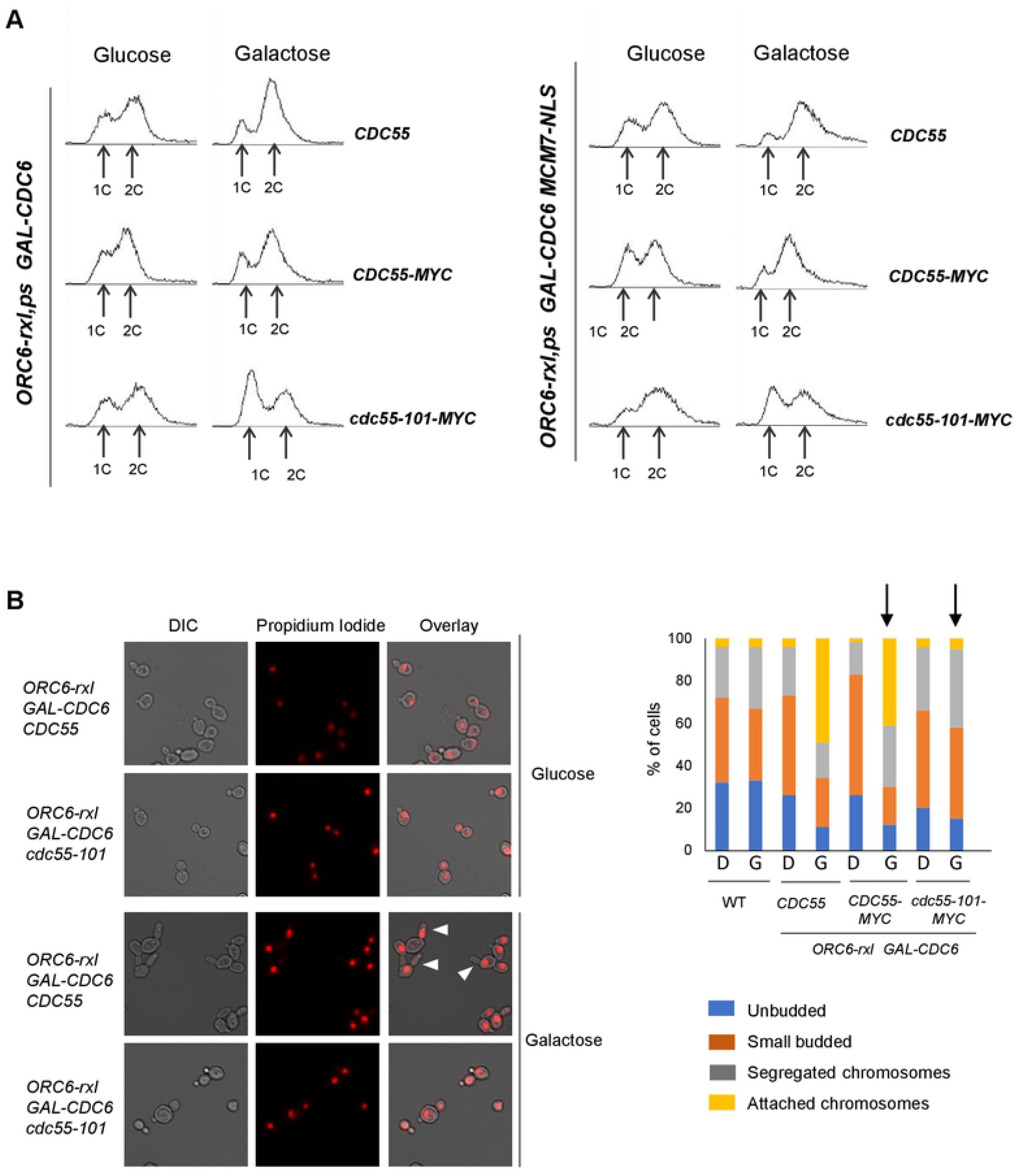
*cdc55-101* bypasses metaphase arrest in re-replicating cells. (A) Cells were grown in raffinose medium were treated with either glucose or galactose for two hours. Cells were fixed and treated with propidium iodide and the cell cycle profile was examined by flow cytometry. (B) Samples from (A) were examined by fluorescence microscopy. 100 cells per genotype and treatment were counted and categorized based on cell cycle stage and chromatid segregation status. Blue=G1, Orange= S/G2, Gray=Mitosis with segregated chromatids, Yellow= Mitosis with attached chromosomes. D is glucose and G is galactose.

Next, we examined chromosome segregation status in *ORC6-rxl GAL-CDC6* cells by propidium iodide staining and fluorescence microscopy. *ORC6-rxl GAL-CDC6* cells showed elongated bud morphology when incubated in galactose as reported previously (Figure 2B, white arrows) [10]. The elongated bud morphology in *ORC6-rxl GAL-CDC6* cells was bypassed when Cdc55 was excluded from nucleus in *cdc55-101* cells (Figure 2B). We next counted the cells based on chromosome segregation status. When incubated in galactose, 41% of *ORC6-rxl GAL-CDC6* cells contained attached chromosomes compared to only 5% with the addition of *cdc55-101* (Figure 2B right, indicated by black arrows). It suggests that mitotic arrest was bypassed when CDC55 was excluded from nucleus (Figure 2B). Taken together, these findings show that nuclear Cdc55 inhibits chromosome segregation in DNA re-replicating cells which leads to metaphase arrest.

Spindle Assembly Checkpoint (SAC) monitors spindle attachment and tension to the kinetochore [17]. Mad2 is a key component of SAC that inhibits the APC and its activator Cdc20, therefore, the cell cycle is arrested in metaphase after SAC activation [18, 19]. It has been previously shown that PP2A-Cdc55 plays a role through SAC activation when the spindle is not attached to the kinetochore [13]. To test if SAC is involved in the metaphase arrest induced upon the response to DNA re-replication, we tested the cell viability of re-replicating cells in the presence or absence of Mad2 with a serial dilution assay. *ORC6-rxl GAL-CDC6* cells showed moderate lethality on galactose plates as previously reported [10] (Figure 3A). Deletion of Mad2 did not affect the cell viability indicating that SAC is dispensable for a re-replication response (Figure 3A). Next, we examined the chromosome segregation status in these cells under fluorescence microscopy. We found that 52% of *ORC6-rxl GAL-CDC6* cells showed attached chromosomes when grown in galactose (Figure 3B, bottom). Similarly, 41% of *ORC6-rxl GAL-CDC6 mad2Δ* cells showed attached chromosomes, indicating that Mad2 does not pay a role in metaphase arrest (Figure 3B, bottom). Furthermore, both *ORC6-rxl GAL-CDC6* cells, with or without *mad2Δ,* showed elongated cell morphology indicative of mitotic arrest (Figure 3B, top with arrows).

**Figure 3.**
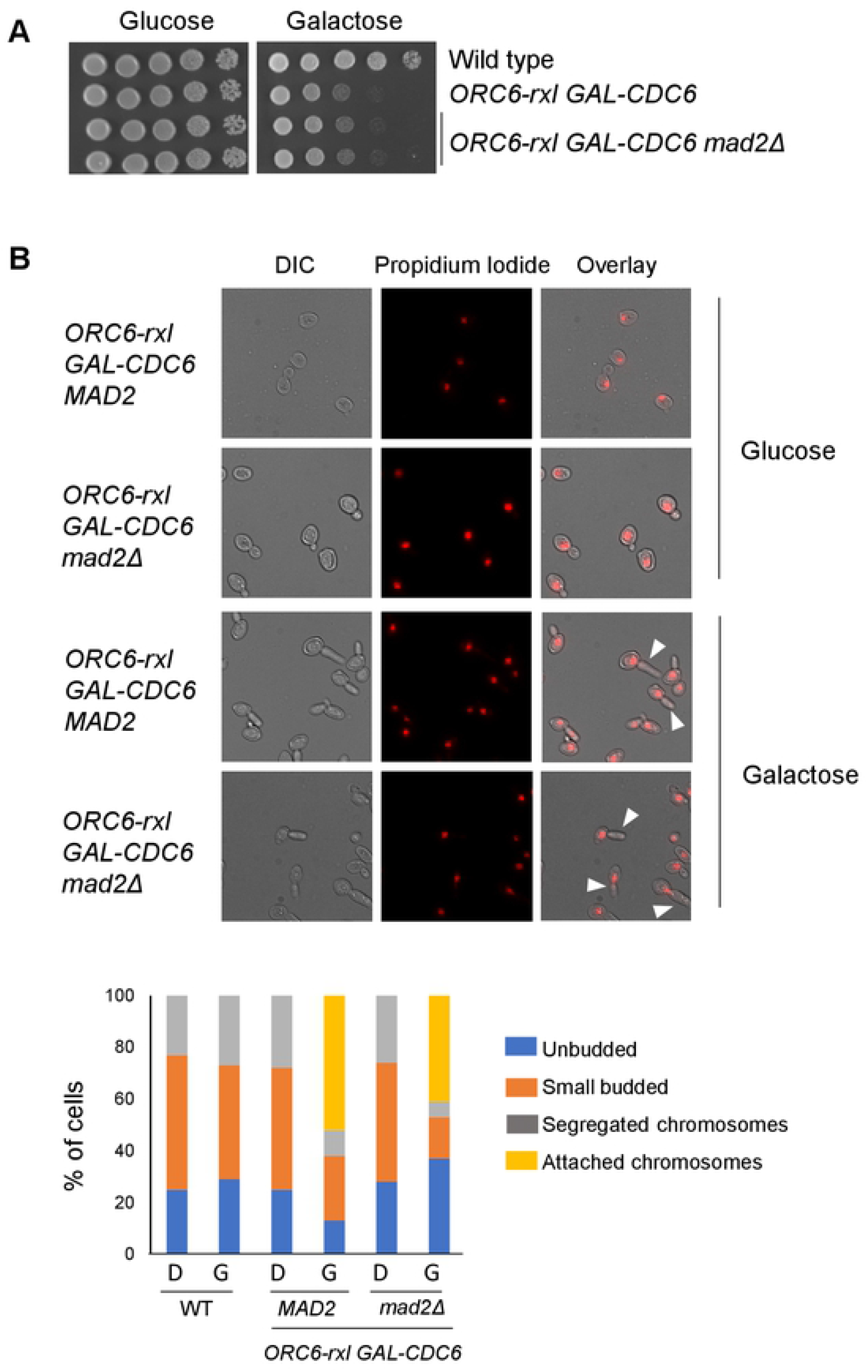
Mad2 does not affect the cell viability of re-replicating cells. (A) Indicated strains were serially diluted 10-fold and spotted on either Glucose- or Galactose-containing plates. (B) (Top) Cells were grown in raffinose-containing medium, and glucose or galactose was added for 2 hours. Samples were fixed and stained with propidium iodide. Cells were examined by fluorescence microscopy. Arrows indicate cells with elongated buds. (Bottom) 100 cells per sample were counted and categorized based on cell cycle stage. Blue=unbudded, Orange=small budded, Gray=large budded cells with segregated chromatids, Yellow=Large budded cells with unsegregated chromatids.

**Figure 4.**
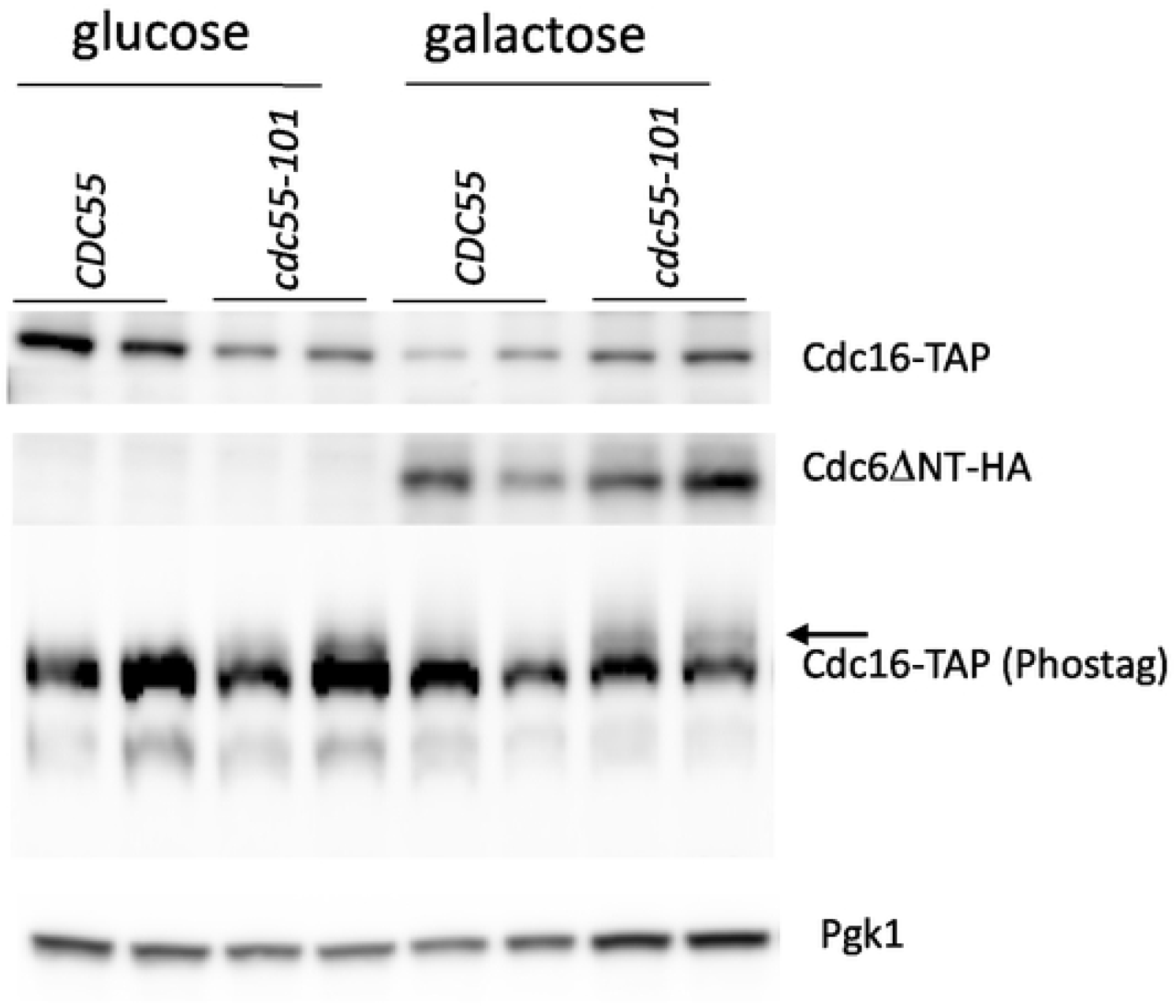
PP2A-Cdc55 dephosphorylates Cdc16 in re-replicating cells. *ORC6-rxl,ps GAL-CDC6DNT CDC16-TAP CDC55 (CDC55*) and *ORC6-rxl,ps GAL-CDC6DNT CDC16-TAP cdc55-101 (cdc55-101*) cells were grown in raffinose medium first, then galactose was added to the media for 4 hours. Samples were collected to detect Cdc16-TAP by western blotting (top) and Cdc16 phosphorylation by Phos-tag analysis (bottom). Pgk1 is shown as a loading control. Two individual clones for each genotype were tested.

It has been shown that PP2A-Cdc55 dephosphorylates and inhibits APC core components such as Cdc16, Cdc23 and Cdc27 [13, 14]. We determined if PP2A-dependent APC inhibition is mediated through Cdc16 dephosphorylation in re-replicating cells. By Phos-tag assay, we examined Cdc16 phosphorylation status in *ORC6-rxl,ps GAL-CDC6ΔNT* cells with or without *cdc55-101. ORC6-rxl,ps GAL-CDC6ΔNT* cells showed hypophosphorylated form of Cdc16 in galactose. Cdc16 was hyperphosphorylated in the presence of *cdc55-101,* showing that PP2A-Cdc55 is responsible for mitotic arrest through Cdc16 dephosphorylation and APC inhibition.

DNA re-replication triggers a DNA damage response which recruits DNA damage repair protein, Rad52 to form foci on double strand DNA breaks [10, 12, 20]. We also tested a possibility that PP2A-Cdc55 might be involved in the DNA damage repair process. Rad52-YFP foci was counted in *ORC6-rxl GAL-CDC6* cells with or without *cdc55-101.* 29.5% of *ORC6-rxl GAL-CDC6* cells formed Rad52-YFP foci in galactose, confirming the DNA damage induction (Figure 5, top). Similarly, 35% of *ORC6-rxl GAL-CDC6 cdc55-101* cells showed Rad52-YFP foci (Figure 5, bottom). Therefore, PP2A-Cdc55 is not involved in the DNA damage repair process.

**Figure 5.**
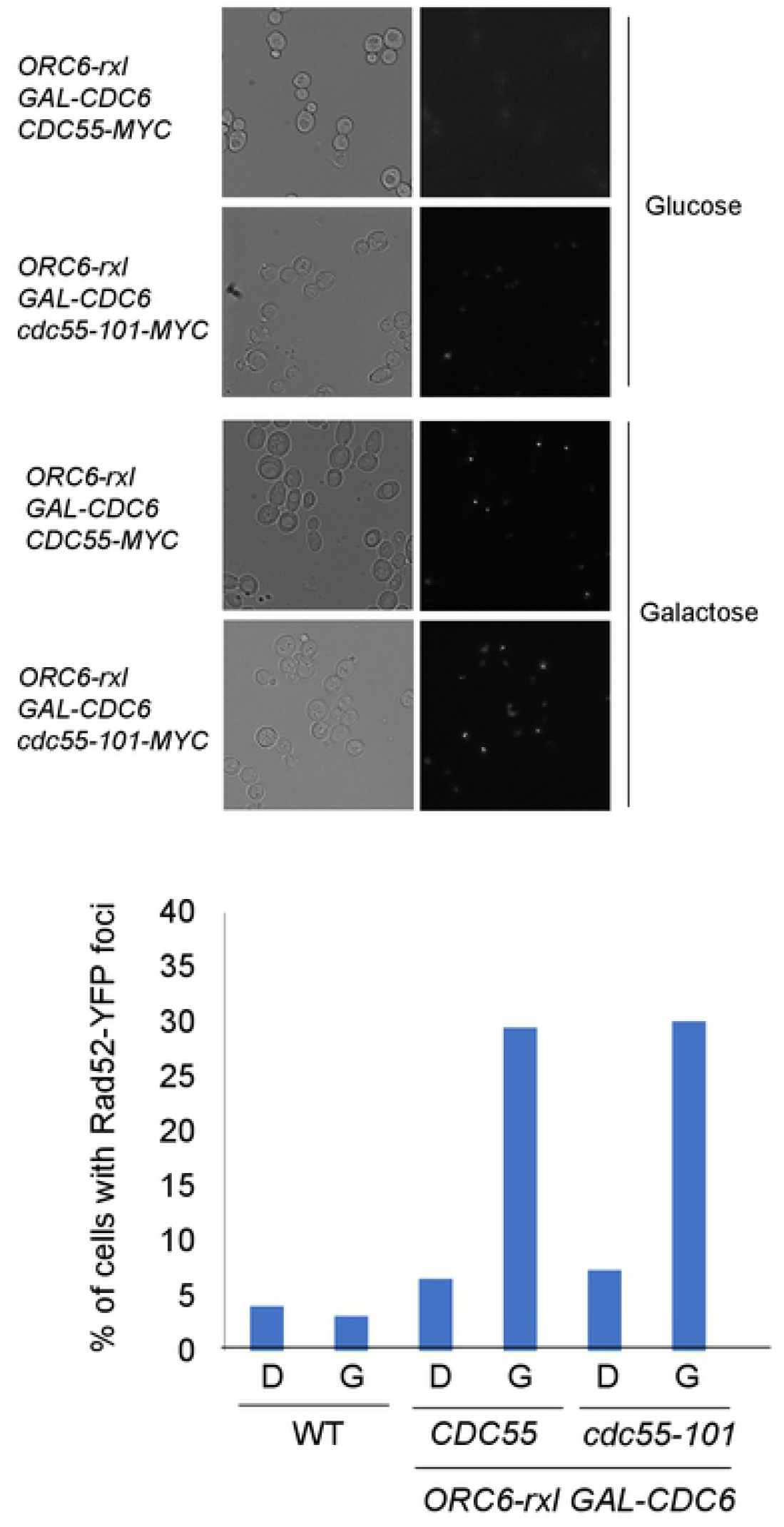
DNA damage induction is not affected by Cdc55. (top) Indicated strains were grown in raffinose-containing medium and then incubated with glucose or galactose for 2 hours. Rad52-YFP foci was counted by fluorescence microscopy. (bottom) Percentage of Rad52-YFP was counted using the samples in A.

Taken together, our data supports a role for PP2A-Cdc55 in mitotic arrest when DNA is over-replicated. This is the first study to show that re-replicating cells arrest the cell cycle in metaphase with attached chromosomes which is dependent on PP2A-Cdc55. Deletion of Cdc55 causes sensitivity to hydroxyurea, replication inhibitor [21]. We recently reported that PP2A-Cdc55 dephosphorylates Pds1, securin, to inhibit spindle elongation during replication stress induced by hydroxyurea. PP2A-Cdc55 may act as a sensor to detect a degree of DNA replication, both in under- and over-replicated DNA. It has been shown that there is a genetic interaction between Cdc55 and Cdc6 [22]. It is of interest to study if PP2A-Cdc55 targets Cdc6 to alter the function. It has been shown that PP2A function is controlled by its localization during an unperturbed cell cycle [13]. The nuclear Cdc55 inhibits anaphase and mitotic exit through the APC and Cdc14 inhibition, respectively. Therefore, Cdc55 nuclear localization might be facilitated by DNA re-replication followed by DNA damage followed by APC inhibition.

## Materials and Methods

### Yeast strains and cell culture

Standard methods of mating and tetrad dissections were used to create strains. All strains are congenic to W303 background. Yeast extract peptone media containing glucose (YEPD) was used to grow cells at 30°C. Serial dilution was performed on YEPD (glucose) or YEPG (galactose) plates. For FACS analysis, cells were first incubated in raffinose-containing media, and then glucose or galactose was added to the media for 2 hours. Synthetic Complete media was used for Rad52-YFP imaging to reduce background signal.

### FACS analysis

Cells were fixed with 70% EtOH and stained with propidium iodide as described before [23]. Flow cytometry analysis was performed using a BD Accuri C6 flow cytometer (BD Biosciences, San Jose, CA). 20,000 cells were analyzed per sample.

### Microscope

Images were taken with a Nikon Eclipse 90i fluorescence microscope using a 60×/1.45 numerical aperture Plan Apochromatic objective lens (Nikon, Tokyo, Japan) equipped with an Intensilight Ultra High Pressure 130-W mercury lamp (Nikon, Tokyo, Japan). Images were captured with a Clara interline charge-coupled device camera (Andor, Belfast, United Kingdom). The images were first obtained with NIS-Elements software (Nikon, Tokyo, Japan), then processed with ImageJ (NIH, Bethesda, MD).

### Western blotting and phos-tag assay

Cells were lysed by agitation in SDS sample buffer with glass beads using FastPrep (MP Biomedicals, OH) for 20 seconds, twice, at speed 6. Protein was separated by SDS-PAGE with 10% polyacrylamide gel. Immuno blotting was performed using anti-HA antibody 3F10 (Roche, IN) at 1:2000 dilution, and anti-Pgk1 (Life Technologies, NY) at 1:2000 as a loading control. Phos-tag acrylamide gel was used to detect phospho-Cdc16-HA (Fujifilm Wako Pure Chemical, Japan).

